# SARS-CoV-2-specific T cell memory is long-lasting in the majority of convalsecent COVID-19 individuals

**DOI:** 10.1101/2020.11.15.383463

**Authors:** Ziwei Li, Jing Liu, Hui Deng, Xuecheng Yang, Hua Wang, Xuemei Feng, Gennadiy Zelinskyy, Mirko Trilling, Kathrin Sutter, Mengji Lu, Ulf Dittmer, Baoju Wang, Dongliang Yang, Xin Zheng, Jia Liu

## Abstract

An unaddressed key question in the current *coronavirus disease 2019* (COVID-19) pandemic is the duration of immunity for which specific T cell responses against the *severe acute respiratory syndrome coronavirus 2* (SARS-CoV-2) are an indispensable element. Being situated in Wuhan where the pandemic initiated enables us to conduct the longest analyses of memory T cell responses against SARS-CoV-2 in COVID-19 convalescent individuals (CIs). Magnitude and breadth of SARS-CoV-2 memory CD4 and CD8 T cell responses were heterogeneous between patients but robust responses could be detected up to 9 months post disease onset in most CIs. Loss of memory CD4 and CD8 T cell responses were observed in only 16.13% and 25.81% of CIs, respectively. Thus, the overall magnitude and breadth of memory CD4 and CD8 T cell responses were quite stable and not inversely correlated with the time from disease onset. Interestingly, the only significant decrease in the response was found for memory CD4 T cells in the first 6-month post COVID-19 disease onset. Longitudinal analyses revealed that the kinetics of SARS-CoV-2 memory CD4 and CD8 T cell responses were quite heterogenous between patients. Loss of memory CD4 T cell responses was observed more frequently in asymptomatic cases than after symptomatic COVID-19. Interestingly, the few CIs in which SARS-CoV-2-specific IgG responses disappeared showed more durable memory CD4 T cell responses than CIs who remained IgG-positive for month. Collectively, we provide the first comprehensive characterization of the long-term memory T cell response in CIs, suggesting that SARS-CoV-2-specific T cell immunity is long-lasting in the majority of individuals.

## Introduction

Antigen-specific T and B cell responses play fundamental roles in the clearance of most viral infections. Additionally, the establishment of T and B cell memory after recovery is essential for protecting the host against disease upon re-exposure. Faced by the unprecedented medical and socioeconomic crisis caused by severe acute respiratory syndrome coronavirus 2 (SARS-CoV-2) and the associated coronavirus disease 2019 (COVID-19), the scientific community has ignited tremendous efforts to map correlates of protection and determinants of immunity against SARS-CoV-2. While antibody-based immunity is relatively well-studied, increasing evidences suggest that T cells may play a fundamental role in the resolution of COVID-19 ^1,2^. The current dogma is that SARS-CoV-2-specific CD4 and CD8 T cell responses, responding at variably high frequencies recognizing multiple epitopes across the viral proteome, can be detected in most individuals both during acute COVID-19 and convalescence afterwards ^3–8^. The magnitude of SARS-CoV-2-specific T cell responses during the early phase is assumed to correlate with the magnitude of antibody responses, and more severe and protracted disease usually drives a more vigorous and, in terms of epitope coverage, broader T cell response ^5,7,8^. However, it has also been observed that cellular and humoral immune responses can become uncoupled in some SARS-CoV-2-exposed individuals, who showed strong specific T cell immunity but lack detectable antibody responses ^9^. It is assumed that this results from antibody responses waning more quickly than T cell responses ^10^ and that SARS-CoV-2-specific antibody responses are rather short-lived, while T cell memory seems to be more durable ^10,11^. However, all available data on analyzing T cell memory were mainly generated from individuals recovering from COVID-19 during a relatively short follow-up period the longest observation duration being less than 60 days post disease onset (dpdo) ^5^. To our knowledge, it is not yet known whether natural infections with SARS-CoV-2 generate long-lasting memory T cell responses and how memory T cell responses changes in a long-term post recovery.

Wuhan was the very first city hit by SARS-CoV-2. Accordingly, all patients who experienced the longest phase of convalescence following COVID-19 reside here or closeby. Wuhan also performed a thorough SARS-CoV-2 RNA test for every resident in May, 2020 to preclude the possibility of local spread of the virus ever since. This enabled us to characterize the long-term memory T cell responses in a cohort of COVID-19 convalescent individuals (CIs) with an unprecedented observation time up to 274 dpdo. Our results suggest that robust SARS-CoV-2 memory T cell responses can be detected in the majority of CIs long-term post recovery.

## Methods

### Subjects

Thirty-one convalescent individuals who resolved their SARS-CoV-2 infection and 11 SARS-CoV-2-unexposed individuals (UIs) were recruited at the Department of Infectious Diseases, Union Hospital, Tongji Medical College, Huazhong University of Science and Technology and the Department of Gastroenterology from April to September 2020. The diagnosis of COVID-19 was based on the Guidelines for Diagnosis and Treatment of Corona Virus Disease 2019 issued by the National Health Commission of China (7^th^ edition). Informed written consent was obtained from each patient and the study protocol was approved by the local medical ethics committee of Union Hospital, Tongji Medical College, Huazhong University of Science and Technology in accordance with the guidelines of the Declaration of Helsinki (2020IEC-J-587).

### Preparation of PBMCs

Peripheral blood mononuclear cells (PBMCs) of SARS-CoV-2-unexposed individuals and patients were isolated using Ficoll density gradient centrifugation (DAKEWE Biotech, Beijing) and were rapidly assessed by flow cytometry analysis without intermittent cryo-preservation.

### Analysis of effector T cell responses

Three pools of lyophilized peptides, consisting mainly of 15-mer sequences with 11 amino acids (aa) overlap, either covering the immunodominant sequences of the surface glycoprotein (S) or the complete sequences of the nucleocapsid phosphoprotein (N) or the membrane glycoprotein (M) of SARS-CoV-2 were used for cell stimulation (PepTivator® Peptide Pools, Miltenyi, Germany). On day 1, PBMCs were resuspended in complete medium (RPMI 1640 containing 10% [v/v] fetal calf serum, 100U/ml penicillin, 100μg/ml streptomycin, and 100μM 4-[2-hydroxyethyl]-1-piperazine ethanesulfonic acid [HEPES] buffer), and stimulated with S, N or M peptide pools (10μg/ml) in the presence of anti-CD28 (1μg/ml; BD Biosciences, USA) and recombinant interleukin (IL)-2 (20U/ml; Hoffmann-La Roche, Italy). Cells without peptide stimulation and anti-CD3-stimulated (1μg/ml; BD Biosciences, USA) cells served as negative and positive controls, respectively. Fresh medium containing IL-2 was added on day 4 and 7. On day 10, cells were restimulated for 5 hours with the same peptide pool in the presence of brefeldin A (BD Biosciences, San Diego, CA). Cells were then tested for IFN-γ, IL-2, and TNF-α expression by intracellular cytokine staining. Specific cytokine responses were calculated by subtracting the background activation (the percentage of cytokine positive cells in the negative control) before further analysis. T cell responses were defined as detectable if the frequency in the specifically stimulated culture exceeded the unstimulated control at least twofold (stimulation index > 2). Samples with responseless positive controls were excluded from further analyses.

### Flow cytometry

Surface and intracellular staining for flow cytometry analysis were performed as described previously ^12,13^. For surface staining, cells were incubated with relevant fluorochrome-labeled antibodies for 30 min at 4°C in the dark. For intracellular cytokine staining, cells were fixed and permeabilized using the Intracellular Fixation & Permeabilization Buffer Set (Invitrogen, USA) and stained with FITC-anti-IFN-γ, PE-anti-IL-2 and APC-anti-TNF-α (BD Biosciences, USA). Approximately 100,000 PBMCs were acquired for each sample using a BD FACS Canto II flow cytometer. Data analysis was performed using the FlowJo software V10.0.7 (Tree Star, Ashland, OR, USA). Cell debris and dead cells were excluded from the analysis based on scatter signals and Fixable Viability Dye eFluor 506.

### Statistical Analysis

Statistical analyses were performed using the SPSS statistical software package (version 22.0, SPSS Inc., Chicago, IL, USA). The Shapiro-Wilk method was used to test for normality. Mann-Whitney t-test, Pearson product-moment correlation coefficient and Fisher’s exact test were used where appropriate. All reported P values were two-sided, and a P value less than 0.05 was considered statistically significant.

## Results

### Characteristics of the study cohort

To characterize SARS-CoV-2-specific memory CD4 and CD8 T cell responses in individuals who had recovered from COVID-19, blood samples derived from 31 CIs together with 11 UIs were assessed. The demographic profiles of all individuals are shown in Table 1. The median period between disease onset and blood sampling was 169 days (range: 83 to 274 days). Among all COVID-19 cases, 56.67% (17/31) were hospitalized and 46.67% (14/31) received oxygen inhalation treatment. Leukopenia and lymphopenia were observed in 52.94% (9/17) and 76.47% (13/17) of tested cases, respectively. Increased C-reactive protein and IL-6 levels were apparent in 70.59% (12/17) and 85.71% (12/14) of tested patients, respectively. Abnormal radiological findings suggesting pneumonia were evident in 74.19% (23/31) CIs by chest computed tomography scans (CT). Sixteen CIs (51.61%) had positive RT-PCR results for viral RNA. All patients were confirmed anti-SARS-CoV-2 IgM and IgG seropositive. At the time of last blood sampling, 45.16% (14/31) were IgG single positive and 29.03% (9/31) were IgM and IgG double positive. Besides, 8 CIs who had become IgG seronegative were purposely recruited to study the interdependence of humoral and cellular immunity. The defining criteria for COVID-19 convalescence were as follows: being afebrile for more than 3 days, resolution of respiratory symptoms, substantial improvement of chest CT images, and two consecutive negative RT-qPCR tests for viral RNA in respiratory tract swab samples obtained at least 24 h apart. At the time of blood sampling, all CIs were negative for viral RNA and had no medical conditions related to COVID-19.

**Table 1.** Baseline characteristics of the Chinese cohort.

| Parameter | Unexposed<br>individuals | Convalescent<br>Individuals |
| --- | --- | --- |
| **n** | 11 | 31 |
| **Gender (M/F)** | 3/8 | 3/28 |
| **Age** | 30.5 | 44.1 |
| **Asymptomatic cases %** | / | 19.35% (6/31) |
| **Mild cases %** |  | 61.29% (19/31) |
| **Severe cases %** | / | 19.35% (6/31) |
| **Days from onset** |  | 169 (83-274) |
| **Days from recovery** | / | 151 (42-249) |
| **Clinical parameters** |  |  |
| Fever % | / | 64.52% (20/31) |
| Respiratory symptoms % | / | 58.06% (18/31) |
| Hospitalized % | / | 56.67% (17/31) |
| Oxygen therapy % | / | 46.67% (14/31) |
| **Laboratory parameters** |  |  |
| Leukopenia % | / | 52.94% (9/17) |
| Lymphopenia % | / | 76.47% (13/17) |
| Increased CRP % | / | 70.59% (12/17) |
| Increased ferritin % | / | 40.00% (4/10) |
| Increased LDH % | / | 40.00% (6/15) |
| Abnormal liver function % | / | 53.33% (8/15) |
| Abnormal renal function % | / | 0 (0/15) |
| Increased CK % | / | 20.00% (3/15) |
| Abnormal blood coagulation % | / | 6.67% (1/15) |
| Increased IL-6 % | / | 85.71% (12/14) |
| **CT scan** |  |  |
| Normal % | / | 25.81% (8/31) |
| Viral pneumonia % | / | 74.19% (23/31) |
| **Virological markers** |  |  |
| RNA positive % | / | 51.61% (16/31) |
| IgG single positive % | / | 45.16% (14/31) |
| IgM & IgG positive % | / | 29.03% (9/31) |
| IgG negative % | / | 25.81% (8/31) |

### Characterization of the long-term memory T cell response specific to SARS-CoV-2

PBMCs of UIs and CIs were re-stimulated with 3 panels of overlapping peptides spanning the SARS-CoV-2 proteins S, N, and M, respectively, to determine memory T cell responses ex vivo. We used an intracellular cytokine staining flow cytometry assay (Fig. S1), and the magnitude of the overall cytokine responses [interferon (IFN)-γ, interleukin (IL)-2, and tumor necrosis factor (TNF)-α] for CD4 and CD8 T cells of all participants are shown in Fig. 1a. Besides, the magnitude and breadth (to how many peptide pools T cells responded) of the IFN-γ, IL-2 or TNF-α-positive T cells are also shown individually in Fig. 1b and 1c. Consistent with previous reports ^4,6^, a proportion of T cells weakly responded to SARS-CoV-2 peptides in UIs (both CD4 and CD8 T cells: 27.27%, 3/11), but with a much lower magnitude than those in CIs (Fig. 1a-1c). In general, memory T cell responses considerably varied in breadth and magnitude between individual CIs. The magnitudes of TNF-α responses against S, IFN-γ or TNF-α responses against N, and IFN-γ responses against M of CD4 and CD8 T cells were significantly positively correlated (Fig. 1d and S2). Memory CD4 T cell responses against a single, two or three peptide pools of the different proteins were detected in 6.45% (2/31), 19.35% (6/31), and 58.06% (18/31) of CIs, respectively (Fig. 1e). Memory CD8 T cell responses against a single, two or three peptide pools of the different proteins were detected in 29.03% (9/31), 16.13% (5/31), and 29.03% (9/31) of CIs, respectively (Fig. 1f). Interestingly, 16.13% (5/31 for CD4) and 25.81% (8/31 for CD8) of CIs did not exhibit memory T cell responses against the three viral proteins (Fig. 1e and 1f). There were only 9.68% (3/31) of CIs who showed no any detectable memory T cell responses against the three proteins for both CD4 and CD8 T cells. Taken together, while the vast majority of CIs had clearly measurable T cell responses against SARS-CoV-2, the data also shows substantial individuality in SARS-CoV-2 memory T cell responses.

**Figure 1.**
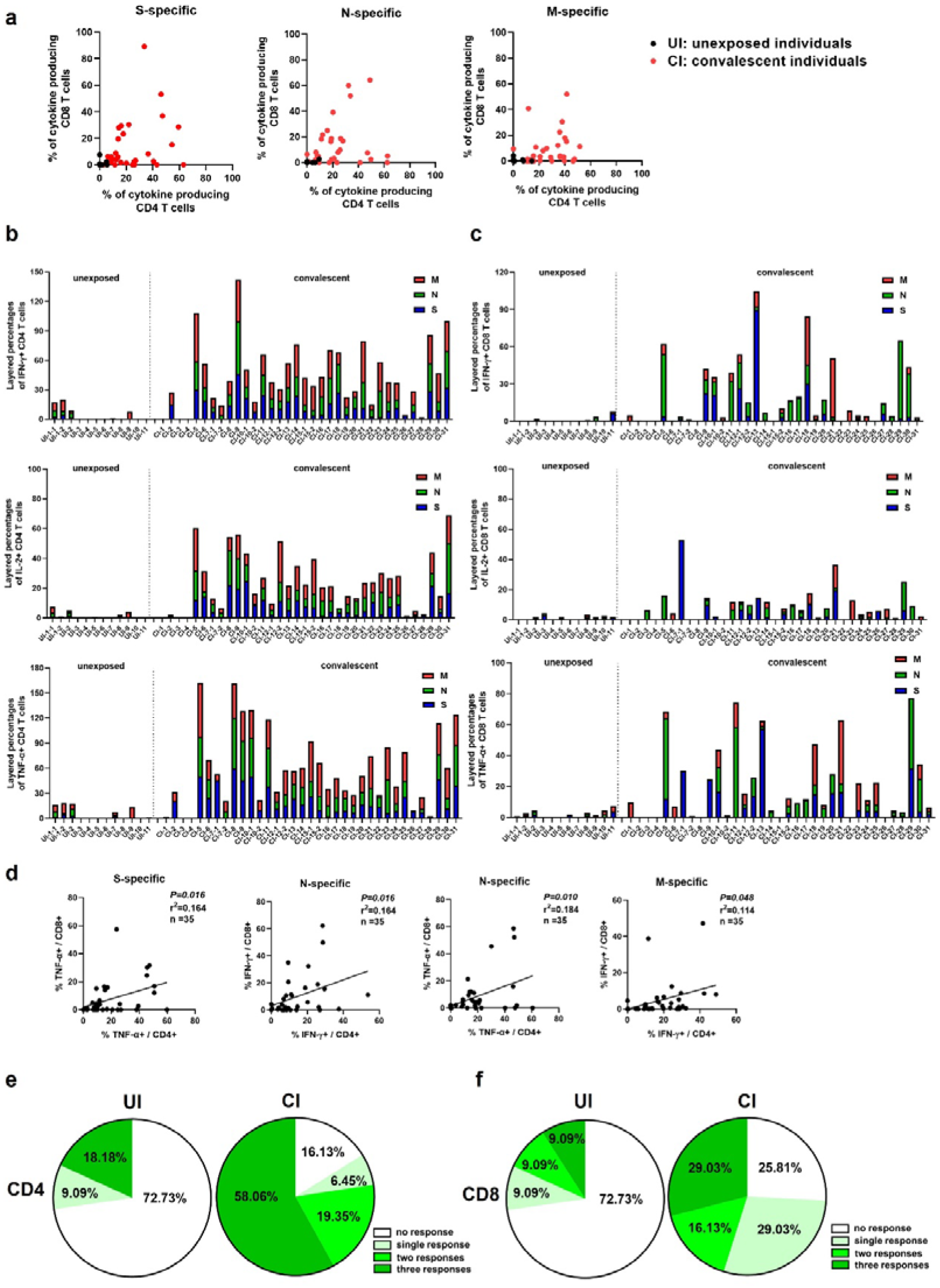
The magnitude and breadth of long-term SARS-CoV-2 memory T cell responses are heterogeneous in COVID-19 convalescent individuals. PBMCs of SARS-CoV-2-unexposed individuals (UI) and COVID-19 convalescent individuals (CI) were tested for responses to 3 panels of overlapping peptides spanning the SARS-CoV-2 S, N, and M, respectively, using intracellular cytokine staining flow cytometry assay. (a) The magnitude of overall cytokine responses of CD4 and CD8 T cells against S, N, and M of SARS-CoV-2 of all participants are shown. (b and c) The magnitude of IFN-γ, IL-2, and TNF-α responses of CD4 and CD8 T cells specific to S, N, and M of SARS-CoV-2 of all participants are also shown individually. Each colored segment represents the source protein corresponding to peptide pools eliciting T cell responses. Bars superimpose percentages of separate T cell culture experiments individually stimulated with indicated antigens. (d) The correlations between the magnitudes of memory CD4 and CD8 T cell responses, as represented by indicated cytokine production, are shown (Pearson product-moment correlation coefficient). (e and f) Breadth of T cell responses of UI and CI. The breadth of T cell responses was calculated by the number of reactive peptide pools of S, N, and M. S: surface glycoprotein; N: nucleocapsid phosphoprotein; M: membrane glycoprotein; IFN: interferon; IL: interleukin; TNF: tumor necrosis factor.

Next, we analyzed the correlation between the magnitude and breadth of the overall SARS-CoV-2 memory T cell responses and the time after disease onset. The CIs were studied up to 9 month after disease onset and we combined the data from all patients for the analysis. In addition, we separately analyzed two different time periods after COVID-19, the first 6 month and the following 3 months for changes in memory T cell responses. For CD4 T cells, the magnitude and breadth of SARS-CoV-2 memory responses against S, N or M showed no significant correlation with days post disease onset (dpdo) (Fig. 2a), suggesting that the CD4 T cell response was relatively stable over time. Interestingly, however, during the first 180 dpdo a significant inverse correlation between the magnitude of the memory CD4 T cell response against S and dpdo was observed (r^2^=0.480, P=0.003, Fig. 2b). In contrast, during the late convalescent phase between 6 and 9 month after COVID-19 the magnitude of memory CD4 T cell responses against S (r^2^=0.327, P=0.041) and N (r^2^=0.328, P=0.041) was positively correlated with dpdo (Fig. 2c). For CD8 T cells, the magnitude and breadth of SARS-CoV-2 memory responses against S, N or M did also not show a significant correlation with dpdo (Fig. 2e). In contrast to CD4 T cells, CD8 T cells did not show a biphasic response during the two different time phases after COVID-19 (Fig. 2d-2f). No significant changes in the magnitude or breath of the CD8 T cell response to any of the SARS-CoV-2 proteins was observed in the early or late phase, with the only exception that a positive correlation between the breadth of memory CD8 T cell responses and dpdo after 180 days was observed (r^2^=0.311, P=0.048, Fig. 2f).

**Figure 2.**
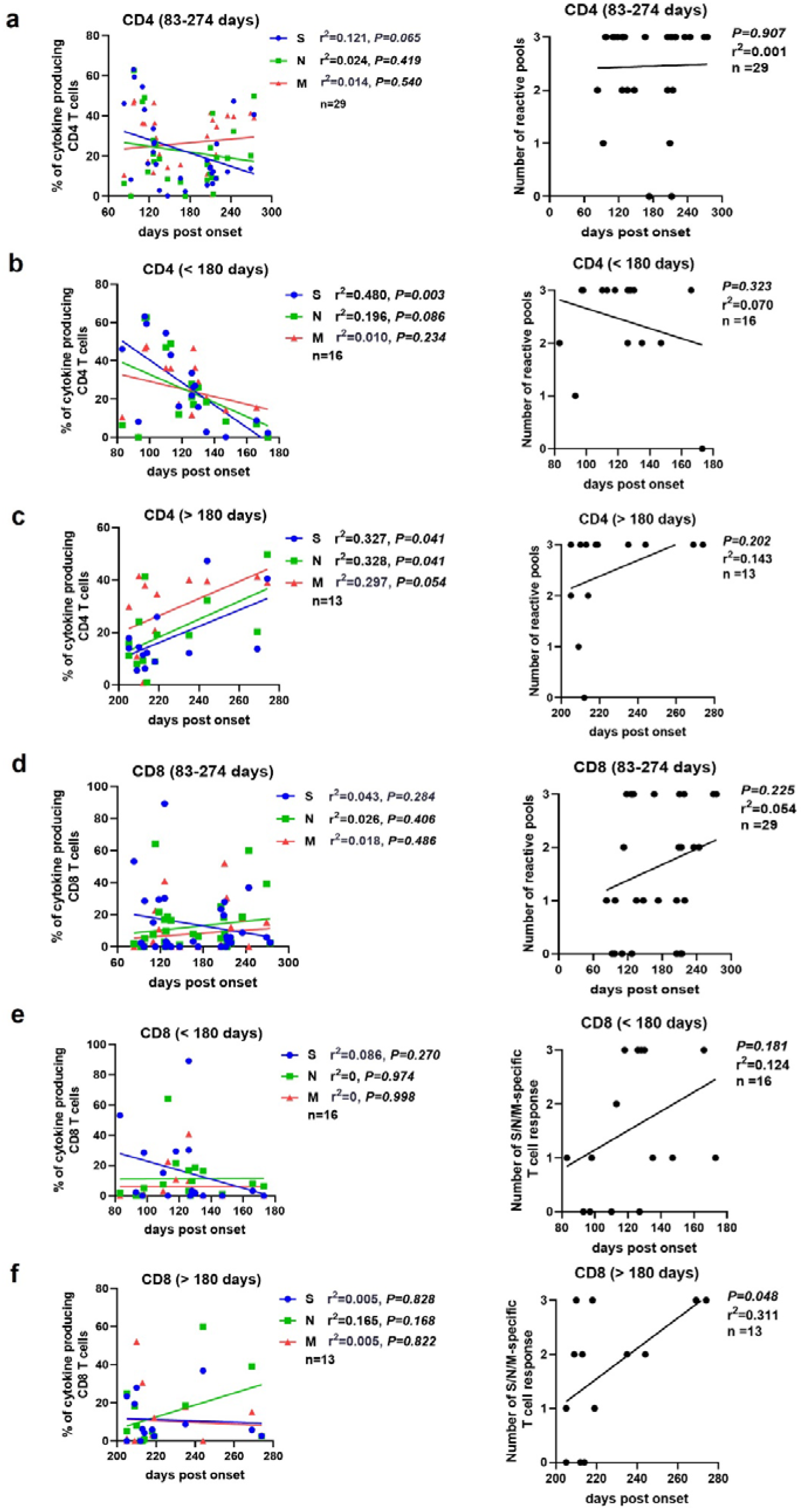
Correlation between the magnitude of SARS-CoV-2 memory T cell responses and the time that had elapsed from disease onset. The correlation between the magnitude of memory CD4 T cell responses specific to S, N and M and days post disease onset up to 274 days (a), within 180 days (b) and over 180 days (c) are shown. The correlation between the magnitude of memory CD8 T cell responses specific to S, N and M and days post disease onset up to 274 days (d), within 180 days (e) and over 180 days (f) are shown. Pearson product-moment correlation coefficient test was used to test the significance and P value and r^2^ value (correlation coefficient) are indicated in each panel. S: surface glycoprotein; N: nucleocapsid phosphoprotein; M: membrane glycoprotein.

These results indicated that the overall SARS-CoV-2 memory CD4 and CD8 T cell responses were long-lasting. However, for memory CD4 T cells a decline in the magnitude of the response was observed during the early recovery phase which was reversed in the following months, highlighting the need for long-term follow up studies such as this.

To further characterize the kinetics of SARS-CoV-2 memory T cell responses, the magnitude of T cell responses were longitudinally examined in more detail in 4 individual CIs. Strong and broad CD4 (in all 4 individuals) and CD8 (3 out of 4 individuals) T cell responses against S, N, and M were detected at the first sampling time point (83-127 dpdo, Fig. 3a-3d). In 2 out of 4 individuals, a decrease in the magnitude of both SARS-CoV-2 memory CD4 and CD8 T cell responses was observed on 147 dpdo and 214 dpdo, respectively (Fig. 3a and 3b), which was most pronounced for the response against the S peptide pool. In contrast, one individual showed sustained SARS-CoV-2 memory CD4 and CD8 T cell responses over time (Fig. 3c), whereas another individual also showed sustained SARS-CoV-2 memory CD4 T cell responses but a strong increase in S-, N-, and M-specific memory CD8 T cell responses, which were undetectable at the early time point in this individual, (Fig. 3d).

**Figure 3.**
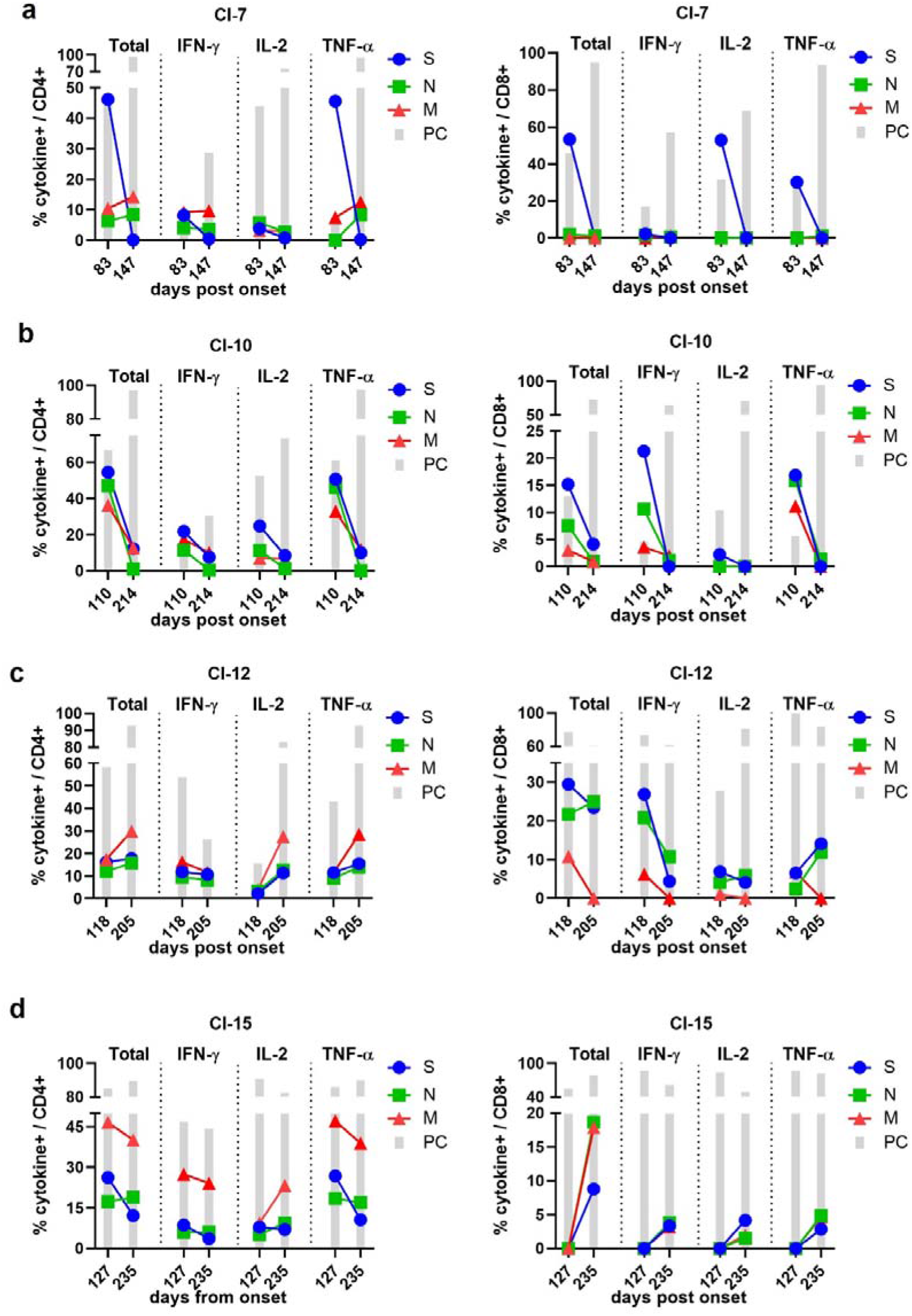
Kinetics of memory T cell responses to SARS-CoV-2 in COVID-19 convalescent individuals. PBMCs were longitudinally collected from 4 COVID-19 convalescent individuals at indicated time points and were tested for memory T cell responses recognizing SARS-CoV-2 S, N or M by using intracellular cytokine staining flow cytometry assay. (a) CI-7; (b) CI-10; (c) CI-12; (d) CI-15. S: surface glycoprotein; N: nucleocapsid phosphoprotein; M: membrane glycoprotein; PC: positive control stimulation; IFN: interferon; IL: interleukin; TNF: tumor necrosis factor.

Taken together, these results suggested that long-term memory T cell responses to SARS-CoV-2 are quite patient-specific and heterogeneous, and may even fluctuate over time in individuals.

### Correlation between the long-term memory T cell response to SARS-CoV-2 and disease severity

Next, we examined the differences in the magnitude and breadth of memory CD4 and CD8 T cell responses in CIs according to their different degrees of COVID-19 severity. CIs were stratified according to the severity of disease into asymptomatic (ACs: 19.35%, 6/31), moderate (MCs: 61.29%, 19/31), and severe COVID-19 cases (SCs: 19.35%, 6/31). No significant difference in the age between the symptomatic and asymptomatic cases was observed. In general, the magnitude of SARS-CoV-2 memory T cell responses against S, N or M, either for the overall or individual cytokine production, were lower in ACs than in MCs and SCs, but the differences were not statistically significant (Fig. 4a, 4b and S3). Also no significant correlations were observed between the magnitude of SARS-CoV-2 memory T cell responses and clinical parameters indicating disease severity, including white blood cell and lymphocyte numbers, IL-6, C-reactive protein, D-dimer, lactate dehydrogenase (LDH), alanine aminotransferase (ALT), aspartate aminotransferase (AST), total bilirubin, serum creatinine, fibrinogen (FIB), and blood urea nitrogen levels (Fig. S4). However, memory CD4 T cell responses against S, N, and M became undetectable in 50% (3/6) of ACs, but only in 5.26% (1/19) of MCs (P=0.031, Fig. 4a). Memory CD8 T cell responses against S, N, and M became undetectable in 50% (3/6) of ACs, but only in 21.05% (4/19) of MCs and 16.67% (1/6) of SCs, respectively (Fig. 4b). No AC showed memory CD8 T cell responses against multiple peptide pools, while 52.63% (10/19) of MCs and 66.67 (4/6) of SCs showed memory CD8 T cell responses to at least 2 different peptide pools (Fig. 4b).

**Figure 4.**
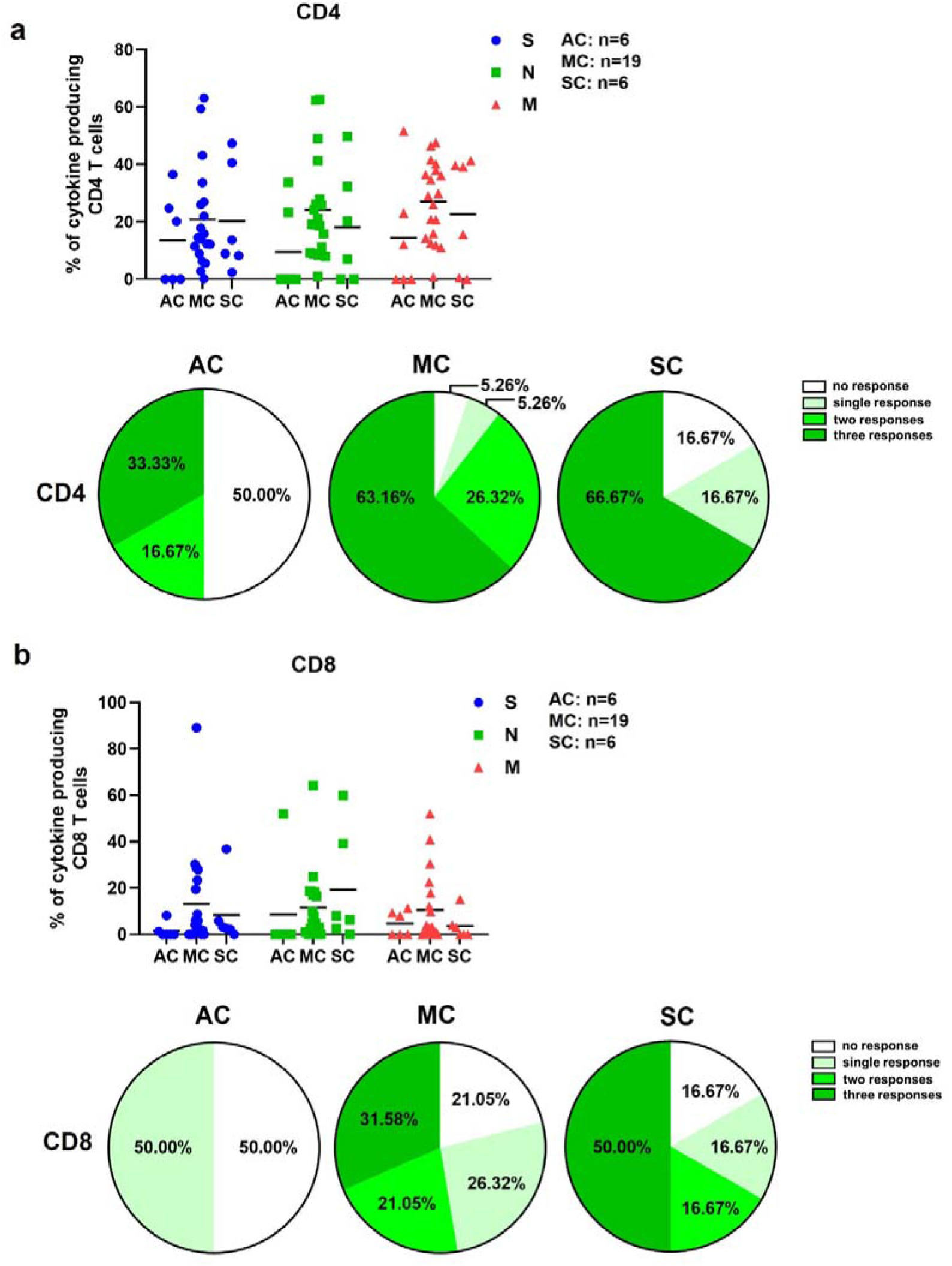
Loss of SARS-CoV-2 memory CD4 T cell responses is more frequent in asymptomatic cases than symptomatic cases. The magnitude and breadth of memory CD4 (a) and CD8 (b) T cell responses are compared between the asymptomatic (AC, n=6), moderate (MC, n=19) and severe (SC, n=6) cases. S: surface glycoprotein; N: nucleocapsid phosphoprotein; M: membrane glycoprotein.

Elderly people are predisposed to develop severe COVID-19 and mortality increases dramatically with age ^14^. We have previously shown that the cytotoxic CD8 T cell response is impaired in elderly COVID-19 patients ^15^. Next, we analyzed correlations between the magnitude and breadth of memory CD4 and CD8 T cell responses and age in CIs. We observed that the breadth, but not the magnitude of memory CD4 T cell responses was inversely correlated with the age of CIs (r^2^=0.162, P=0.016, Fig. 5a). No significant correlation between the magnitude and breadth of memory CD8 T cell responses and the age of CIs were observed (Fig. 5b).

**Figure 5.**
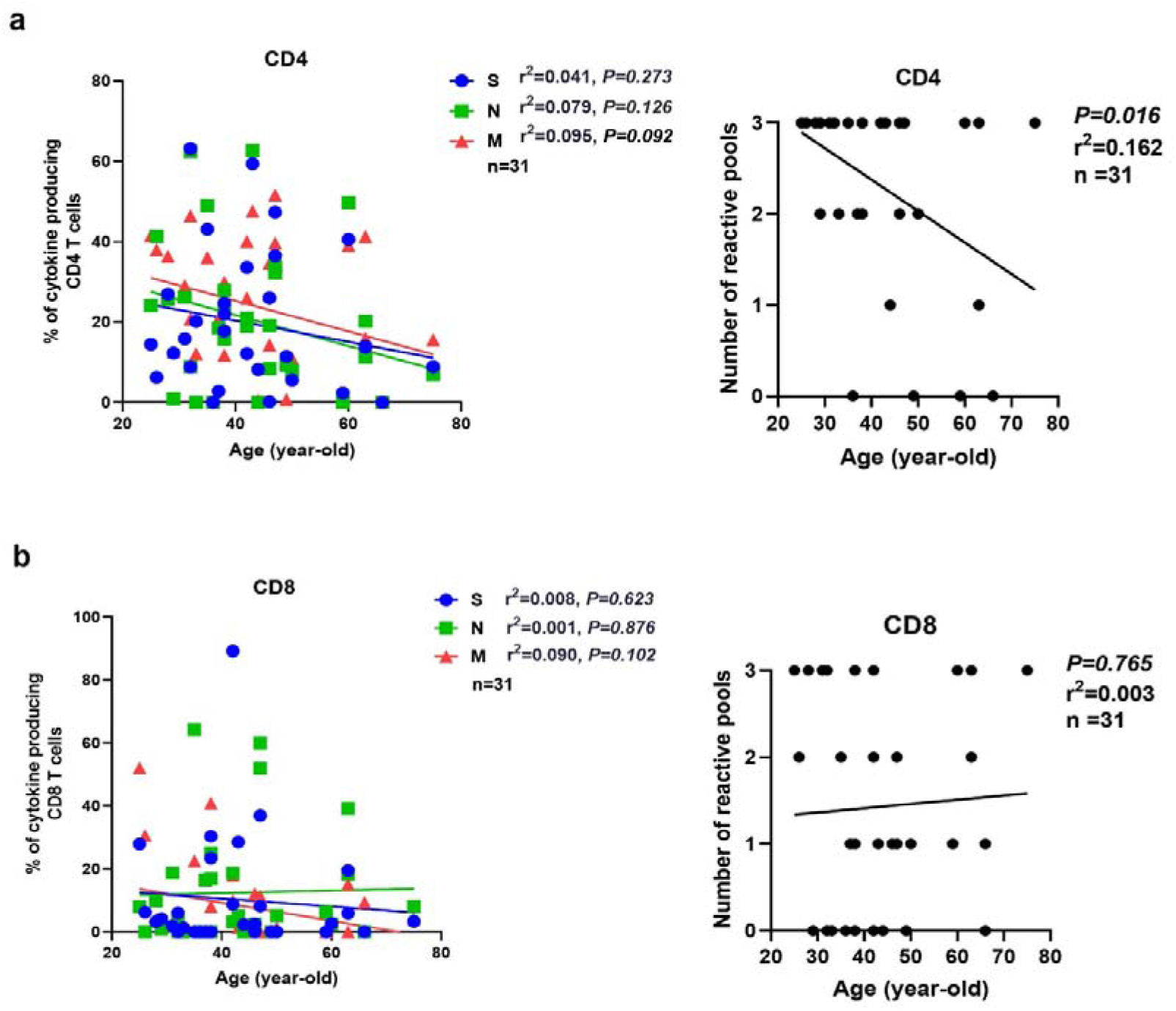
The breadth of long-term SARS-CoV-2 memory CD4 T cell responses is negatively correlated with the age of COVID-19 convalescent individuals. The correlation between the magnitude and breadth of memory CD4 (a) and CD8 (b) T cell responses specific to S, N and M and age are shown. Pearson product-moment correlation coefficient test was used to test the significance and P value and r^2^ value (correlation coefficient) are indicated in each panel. S: surface glycoprotein; N: nucleocapsid phosphoprotein; M: membrane glycoprotein.

### CD4 memory T cell responses in individuals who lost their IgG response to SARS-CoV-2

During the acute phase of COVID-19, T cell responses positively correlated with the magnitude of antibody responses ^5,7,8^. However, to our knowledge it is not clear whether this association is maintained during the long-term convalescence. To this end, we compared memory T cell responses and antibody responses in CIs from 83 to 274 dpdo. As shown in Fig. S5, the magnitude of memory CD4 and CD8 T cell responses against S and N showed no significant correlation with the titers of corresponding IgG against S and N. Form our large convalescent out-patient cohort very few patients lose their SARS-COV-2-specific IgG responses over time. We were interested if those patients still kept their memory T cells. We therefore selected 8 IgG-seronegative CIs and compared them to 23 seropositive CIs. At the time point of last sampling the age of the IgG-seronegative CIs was significantly lower, and the dpdo was significantly higher, than those of the 23 IgG-seropositive CIs (Fig. S6a). To overcome this bias, we compared the magnitude and breadth of memory T cell responses of IgG-seronegative CIs with 7 selected IgG-seropositive CIs with comparable age and dpdo (Fig. S6b). Interestingly, memory CD4 T cell responses against N and M were significantly higher in IgG-seronegative CIs than those in IgG-seropositive CIs (Fig. 6a). A tendency of increased memory CD4 T cell response against S in IgG-seronegative CIs was also observed, although the difference remained close to the borderline of statistical significance (P=0.052, Fig. 6a). All IgG-seronegative CIs showed memory CD4 T cell responses to at least 2 peptide pools, while 28.57% IgG-seropositive CIs showed no memory CD4 T cell responses to S, N or M (Fig. 6b). In contrast to CD4 T cells, no significant differences in the magnitude and breadth of memory CD8 T cell responses between the IgG-seronegative and -seropositive CIs were observed (Fig. 6c and 6d), indicating that CD8 T cells seem to be less correlated with humoral immune responses as compared to CD4 T cells.

**Figure 6.**
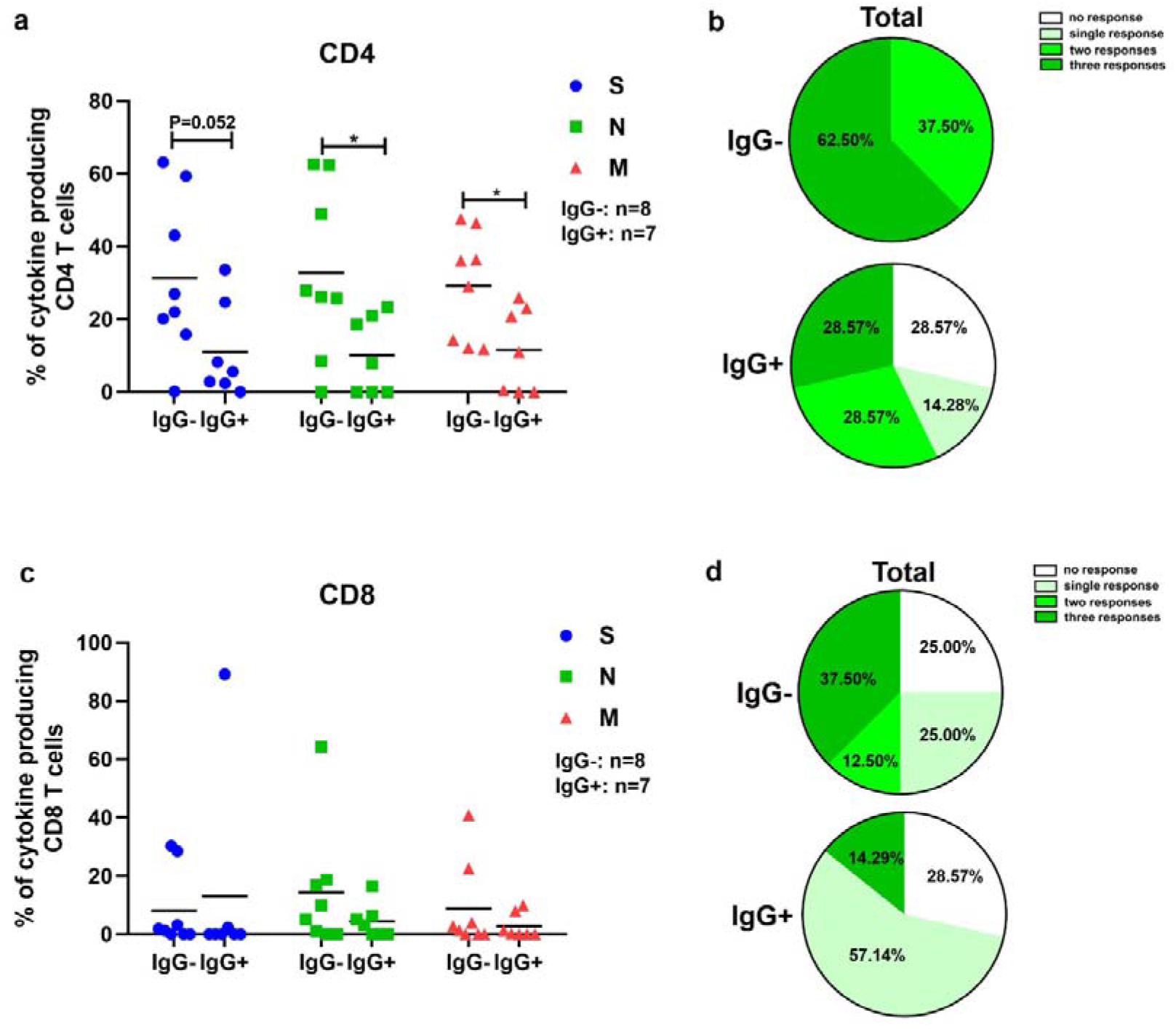
The long-term SARS-CoV-2 memory CD4 T cell responses is robust in IgG-seronegative COVID-19 convalescent individuals. The magnitude (a) and breadth (b) of memory CD4 T cell responses are compared between IgG-seronegative (IgG-, n=8) and IgG-seropositive (IgG+, n=7) CIs. The magnitude (c) and breadth (d) of memory CD8 T cell responses are compared between IgG-seronegative (IgG-, n=8) and IgG-seropositive (IgG+, n=7) CIs. Statistically significant differences are indicated by asterisks (* < 0.05, Non-parametric Mann-Whitney test). S: surface glycoprotein; N: nucleocapsid phosphoprotein; M: membrane glycoprotein.

## Discussion

One of the most important and challenging questions facing medicine today concerns the extent to which immunity develops and persists following COVID-19. Previous studies suggest that the persistence of protective immunity against different coronaviruses varies significantly, since those against seasonal coronavirus are short-lived ^16^ while those against SARS and *middle east respiratory syndrome coronavirus* (MERS) are described to last longer ^6,17,18^. Recent studies have demonstrated that macaques infected with SARS-CoV-2 are resistant to reinfection with the same virus isolate following recovery from their initial infection, suggesting the cellular and/or humoral immunity facilitated by the primary infection might have protected the same nonhuman primates against secondary encounters ^19,20^. However, in both studies, reinfections with SARS-CoV-2 were carried out within a relative short time window (4 and 5 weeks after the primary infection). In contrast to the observation in the macaque model, there are some reports demonstrating the principle possibility of reinfections with SARS-CoV-2 in humans ^21–24^. It has been suggested that the lifespan of the humoral response following SARS-CoV-2 infection is relatively short, especially in mild and asymptomatic cases ^25^. Some believe that although SARS-CoV-2 infection may blunt long-lived antibody responses, immune memory might still be achieved through virus-specific memory T cell responses ^2^, which have been detected in most recently recovered individuals, including asymptomatic cases and those with undetectable antibody responses ^9^. Here we provide, to our knowledge, the first characterization of long-term memory T cell responses in a cohort of COVID-19 convalescent individuals up to 9 months following primary SARS-CoV-2 infection. We show that the magnitude and breadth of long-term memory T cell responses to SARS-CoV-2 are heterogeneous. While the majority of CIs demonstrate strong and broad memory T cell responses up to 9 months post disease onset, some individuals have lost their T cell responses against the studied antigens within half a year. The magnitude of SARS-CoV-2 memory CD4 T cell response is inversely correlated with the time that had elapsed from disease onset within 180 days, suggesting SARS-CoV-2 memory CD4 T cell response may wane over time at the early months following primary SARS-CoV-2 infection. Intriguingly, half of the asymptomatic cases have lost their memory CD4 and CD8 T cell responses, suggesting the memory T cell responses might be less durable in asymptomatic cases than in symptomatic cases. The breadth of memory CD4 T cell responses were inversely correlated with the age of the patients, suggesting the memory T cell responses might also be less durable in elderly individuals. Moreover, the kinetics of memory T cell responses are heterogeneous in the herein examined CIs, while some show a sharp decline of memory T cell responses over time, others show rather sustained or even increasing memory T cell responses. Our data document a durability of cellular immunity against SARS-CoV-2, however, for a fraction of elderly individuals with asymptomatic infections a considerable waning of cellular immunity may occur. Our results also suggest that the intensity of SARS-CoV-2 memory T cell responses detected in peripheral blood may fluctuate over time in CIs, which is unlikely to be caused by reexposion to SARS-CoV-2, since the the possibility of local spread of the virus in Wuhan and nearby area has been precluded by the thorough SARS-CoV-2 RNA test conducted in May for every resident. Future studies are needed to closely monitor the SARS-CoV-2 memory T cell responses to address how the intensities of these responses are regulated in CIs.

Different from the observation during and shortly after the acute phase of SARS-CoV-2 infection ^5,7,8^, we observe that the magnitudes of long-term SARS-CoV-2-specific cellular and humoral responses are not positively correlated with each other. In contrast, IgG-seronegative CIs demonstrate even stronger SARS-CoV-2-specific memory CD4 T cell responses than IgG-seropositive CIs. A recent study started to investigate the possible mechanisms of short-lived antibody responses observed in COVID-19 patients and has reported that germinal centers in secondary lymphoid organs were largely absent during the acute phase of COVID-19 ^26^. The authors speculate that the absence of germinal centers is a result of abundant Th1 cell responses and aberrant extra-follicular TNF-α accumulation ^26^. Consistently, our current observation, that CIs with short-lived antibody responses demonstrate an increased magnitude of SARS-CoV-2-specific CD4 T cell responses, provides the first evidence that the above-mentioned effect may extent to a far longer period in the convalescent phase of COVID-19. Although it remains unclear which arms of the adaptive immune response are responsible for protection against SARS-CoV-2 infection, our data demonstrate that CIs may possess at least one arm of the adaptive immune response against SARS-CoV-2 long-term post recovery. Further characterization of the protective roles as well as the interaction of cellular and humoral immune responses against SARS-CoV-2 has significant implications for vaccine development and application especially in terms of the need for booster vaccinations.

Taken together, we provide the first comprehensive characterization of the long-term memory T cell responses against SARS-CoV-2, suggesting that the SARS-CoV-2-specific T cell immunity is sustained in the majority of CIs up to 9 months post infection. The observation that convalescent individuals turning IgG-seronegative generated robust and sustained memory T cell responses further suggests that natural infection could prevent recurrent episodes of severe COVID-19.

## Acknowledgement

This work is supported by the Fundamental Research Funds for the Central Universities (2020kfyXGYJ028, 2020kfyXGYJ046 and 2020kfyXGYJ016), the National Natural Science Foundation of China (81861138044 and 91742114), the National Science and Technology Major Project (2017ZX10202203), and the Medical Faculty of the University of Duisburg-Essen and Stiftung Universiätsmedizin, University Hospital Essen, Germany. M.T., K.S., M.L., and U.D. receive funding from the Deutsche Forschungsgemeinschaft (DFG) for example through the RTG1949/2.

