## Supplemental Materials for "SARS-CoV-2-specific T cell memory is long-lasting in the majority of convalsecent COVID-19 individuals"

**Supplementary Materials**

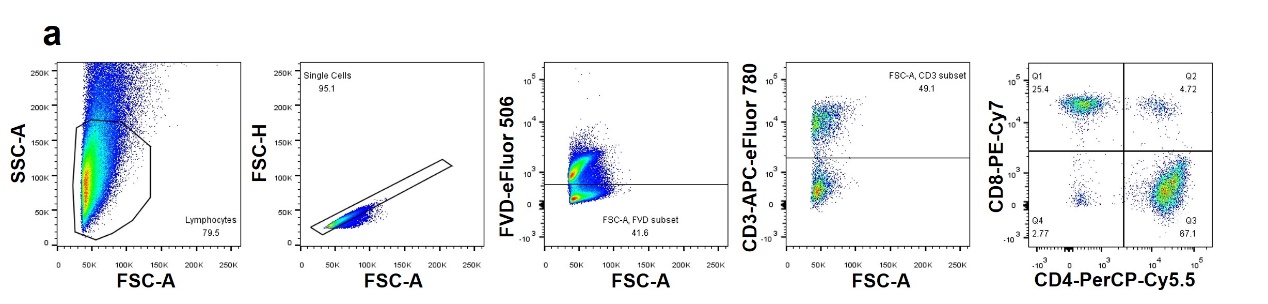

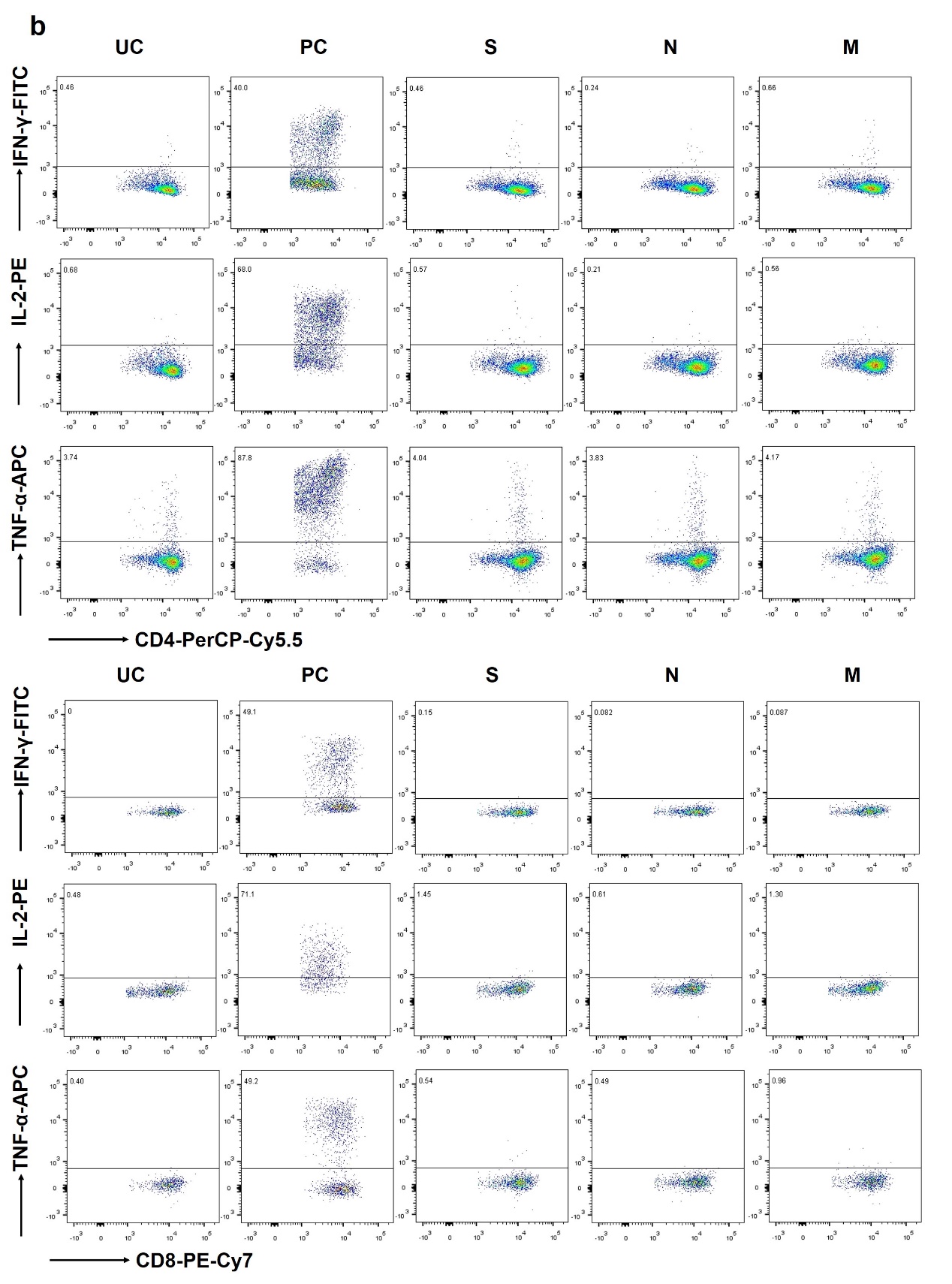

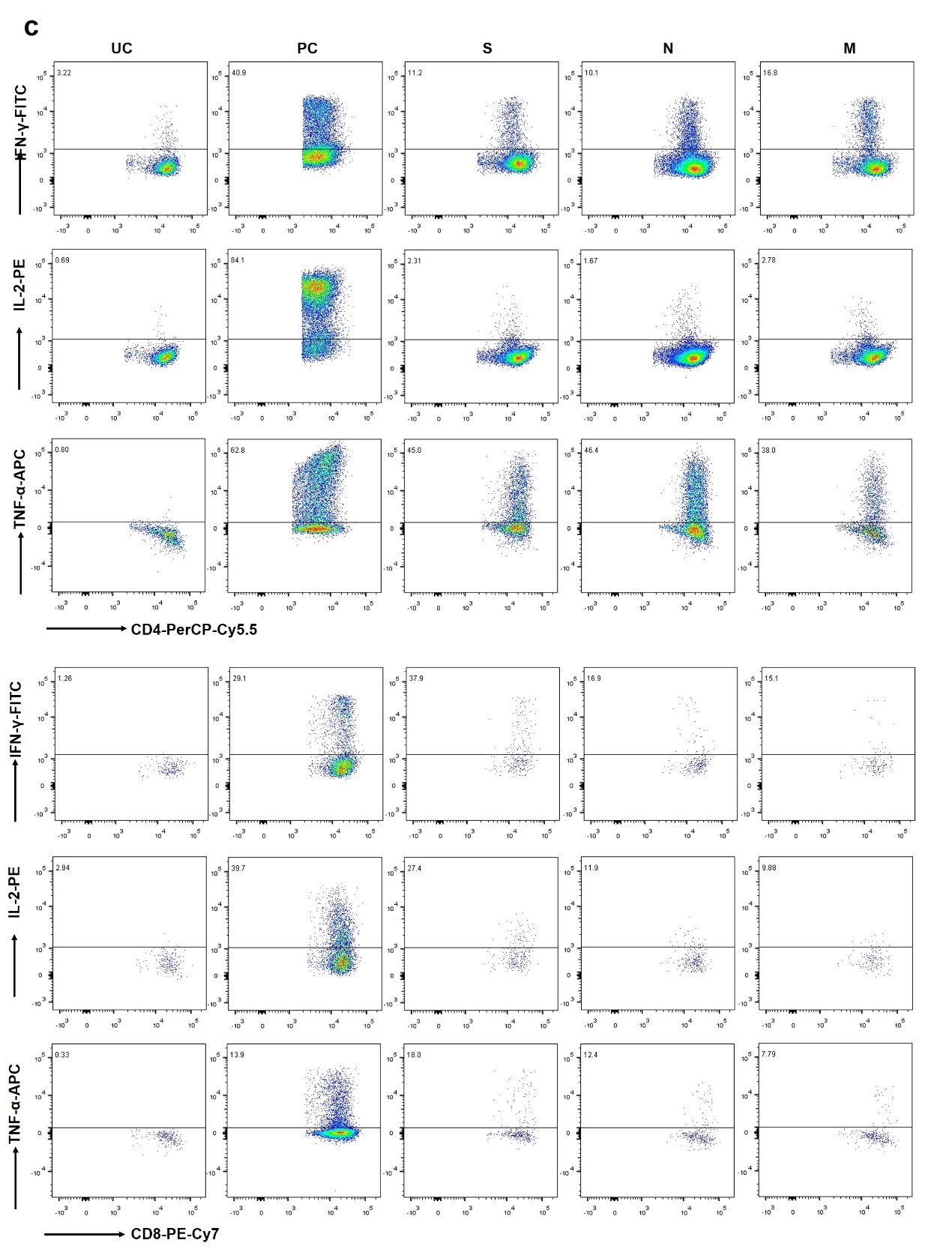

**Figure S1.** **Characterization of memory T cell responses specific to SARS-CoV-2 viral proteins in unexposed individuals and COVID-19 convalescent individuals.** (a) Exemplary strategy for gating CD4 and CD8 T cells by flow cytometry. Exemplary gating strategy for analyzing memory CD4 and CD8 T cell responses specific to SARS-CoV-2 S, N and M in UI (b) and CI (c). UI: SARS-CoV-2-unexposed individuals; CI: COVID-19 convalescent individuals. UC: unstimulated control; PC: positive control stimulation; S: surface glycoprotein; N: nucleocapsid phosphoprotein; M: membrane glycoprotein; IFN: interferon; IL: interleukin; TNF: tumor necrosis factor.

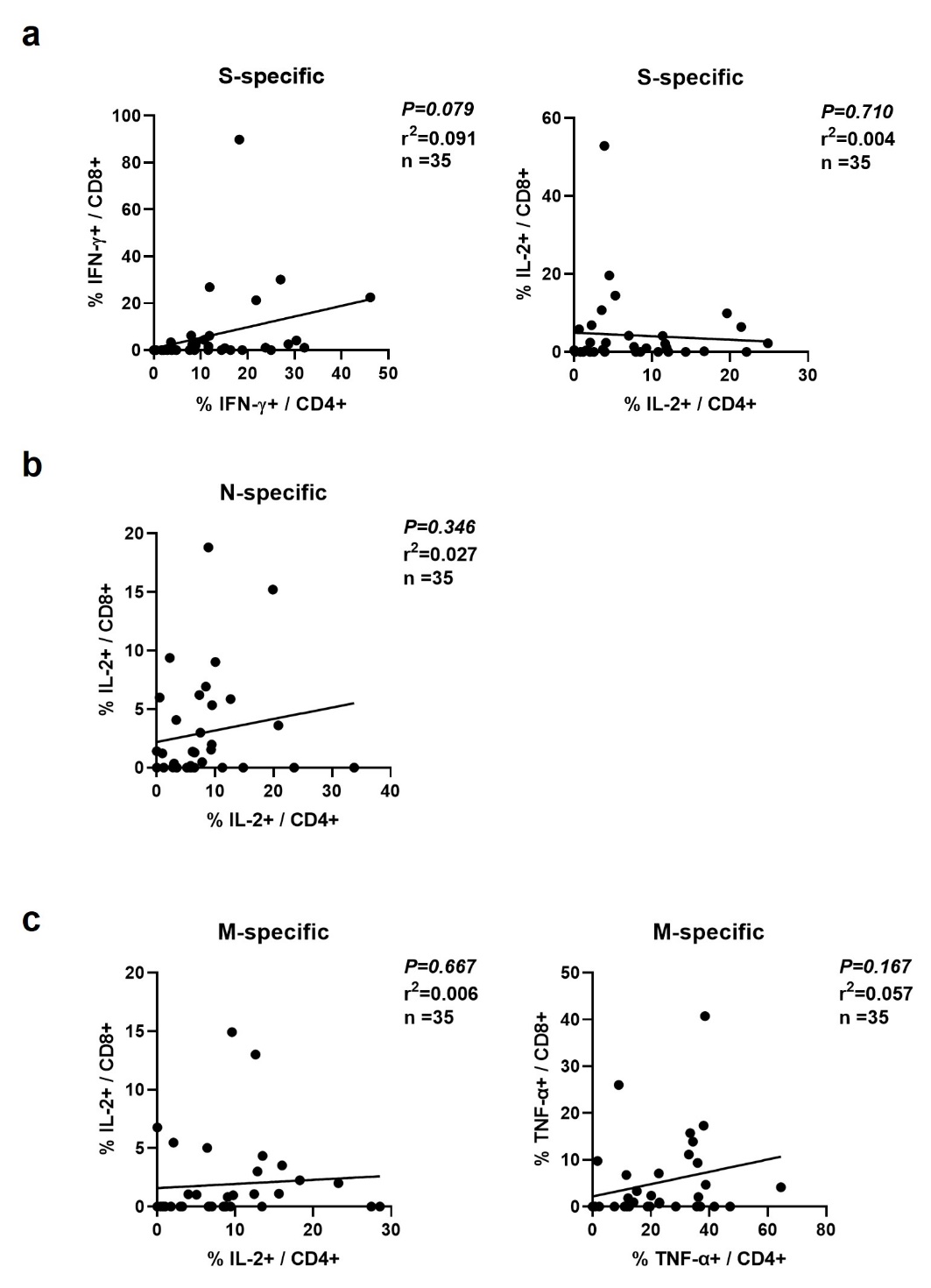

**Figure S2.** **Correlation between the magnitudes of SARS-CoV-2 memory CD4 and CD8 T cell responses.** The correlation between the magnitude of memory CD4 and CD8 T cell responses specific to S (a), N (b), and M (c) are shown. Pearson product-moment correlation coefficient test was used to test the significance and P value and r^2^ value (correlation coefficient) are indicated in each panel. S: surface glycoprotein; N: nucleocapsid phosphoprotein; M: membrane glycoprotein; IFN: interferon; IL: interleukin; TNF: tumor necrosis factor.

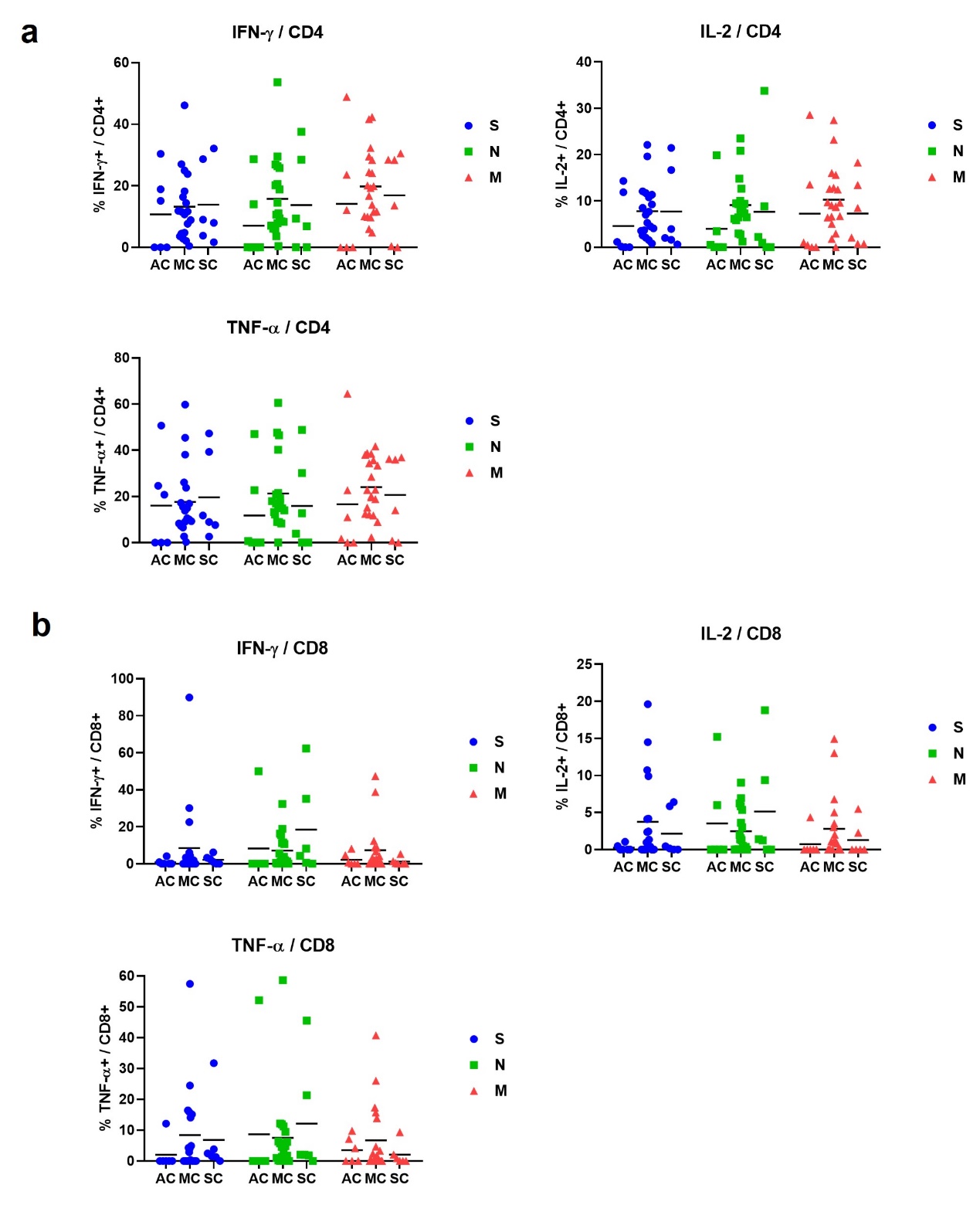

**Figure S3.** **Characterization of memory T cell responses specific to SARS-CoV-2 in COVID-19 convalescent individuals with different disease severity.** The magnitude of memory CD4 (a) and CD8 (b) T cell responses are compared between the asymptomatic (AC, n=6), moderate (MC, n=19) and severe (SC, n=6) cases. S: surface glycoprotein; N: nucleocapsid phosphoprotein; M: membrane glycoprotein; IFN: interferon; IL: interleukin; TNF: tumor necrosis factor.

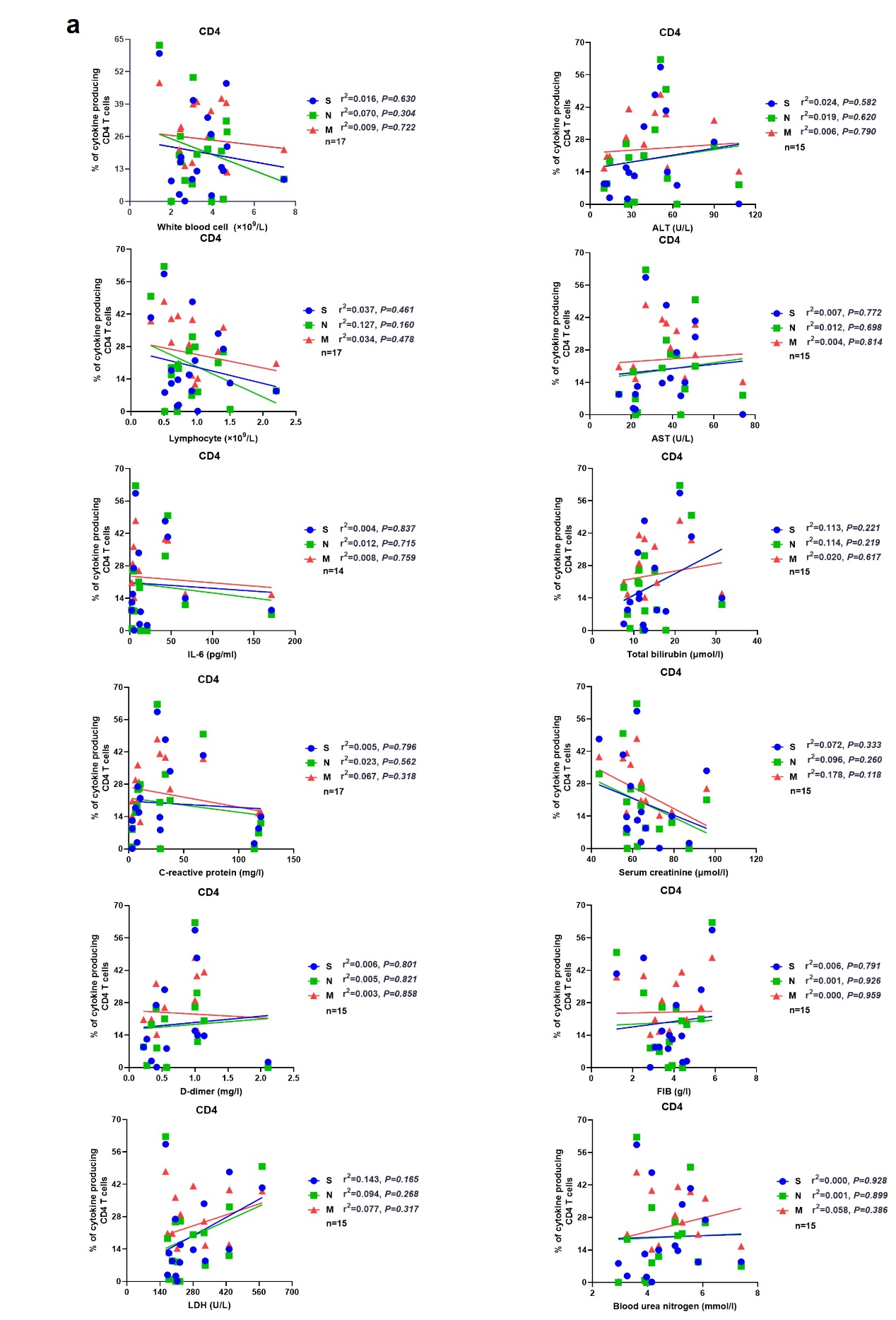

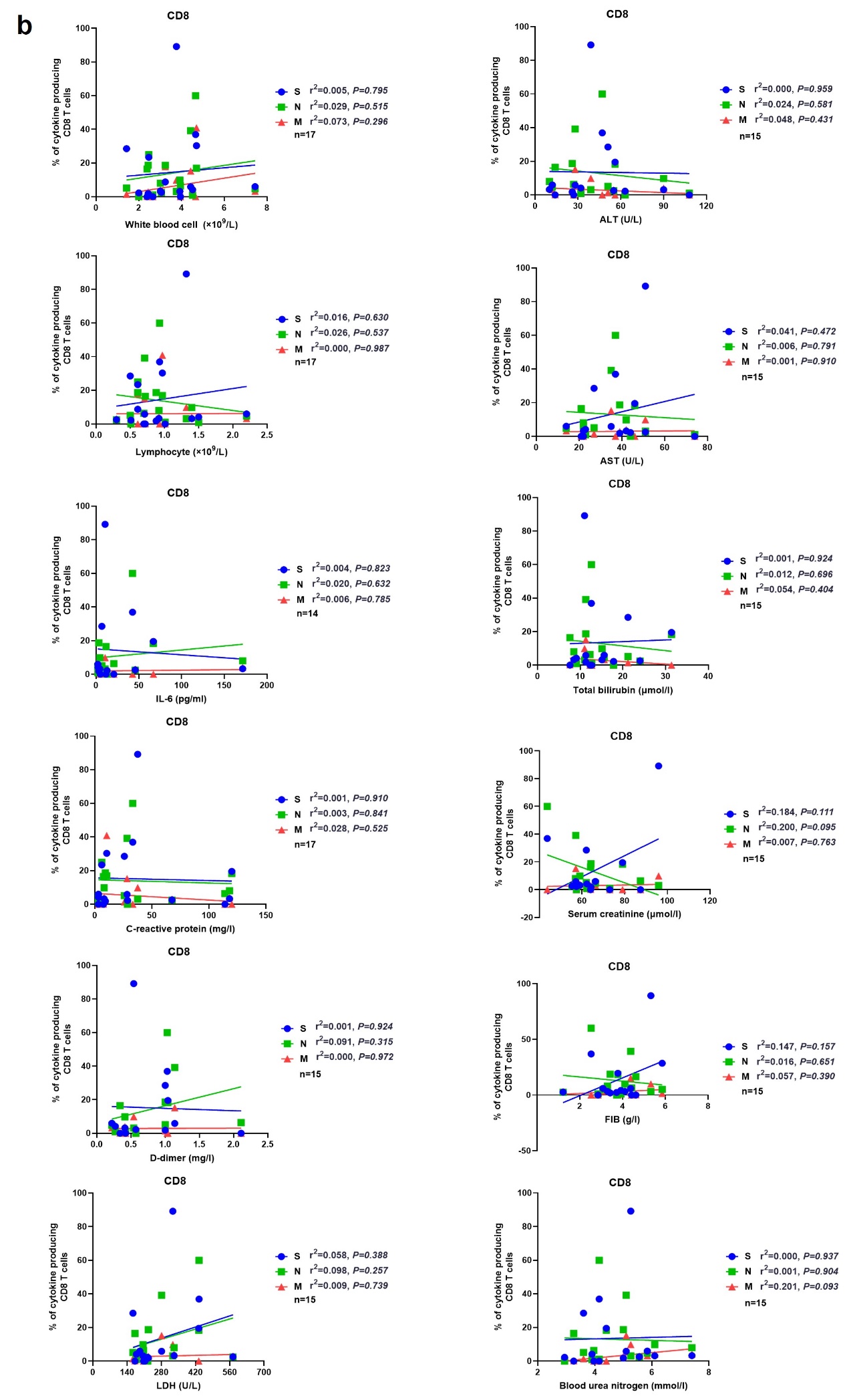

**Figure S4. Correlation between the magnitude of SARS-CoV-2 memory T cell responses and clinical parameters indicating COVID-19 severity.** The correlations between the magnitude of memory CD4 (a) and CD8 (b) T cell responses and white blood cell and lymphocyte numbers, IL-6, C-reactive protein, D-dimer, LDH, ALT, AST, total bilirubin, creatinine, FIB, and blood urea nitrogen levels are shown respectively. Pearson product-moment correlation coefficient test was used to test the significance and P value and r^2^ value (correlation coefficient) are indicated in each panel. S: surface glycoprotein; N: nucleocapsid phosphoprotein; M: membrane glycoprotein; IL: interleukin; LDH: lactate dehydrogenase; ALT: alanine aminotransferase; AST: aspartate aminotransferase; FIB: fibrinogen.

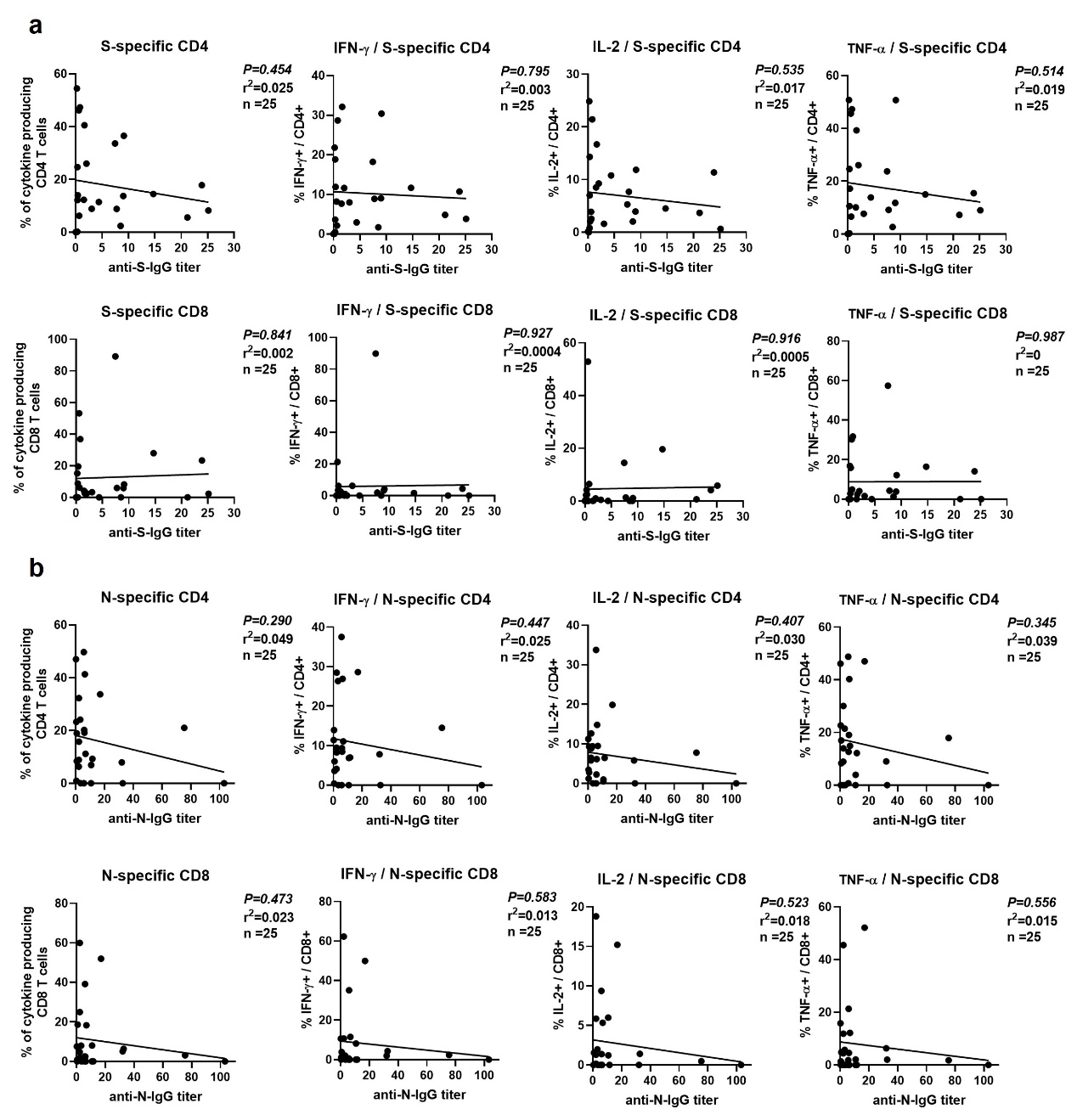

**Figure S5. Correlation between the magnitude of SARS-CoV-2 memory T cell responses and the titers of corresponding IgG antibodies.** The correlations between the magnitude of memory T cell responses specific to S (a) or N (b) and the titers of S-specific IgG or N-specific IgG antibodies are shown respectively. Pearson product-moment correlation coefficient test was used to test the significance and P value and r^2^ value (correlation coefficient) are indicated in each panel. S: surface glycoprotein; N: nucleocapsid phosphoprotein; M: membrane glycoprotein; IFN: interferon; IL: interleukin; TNF: tumor necrosis factor.

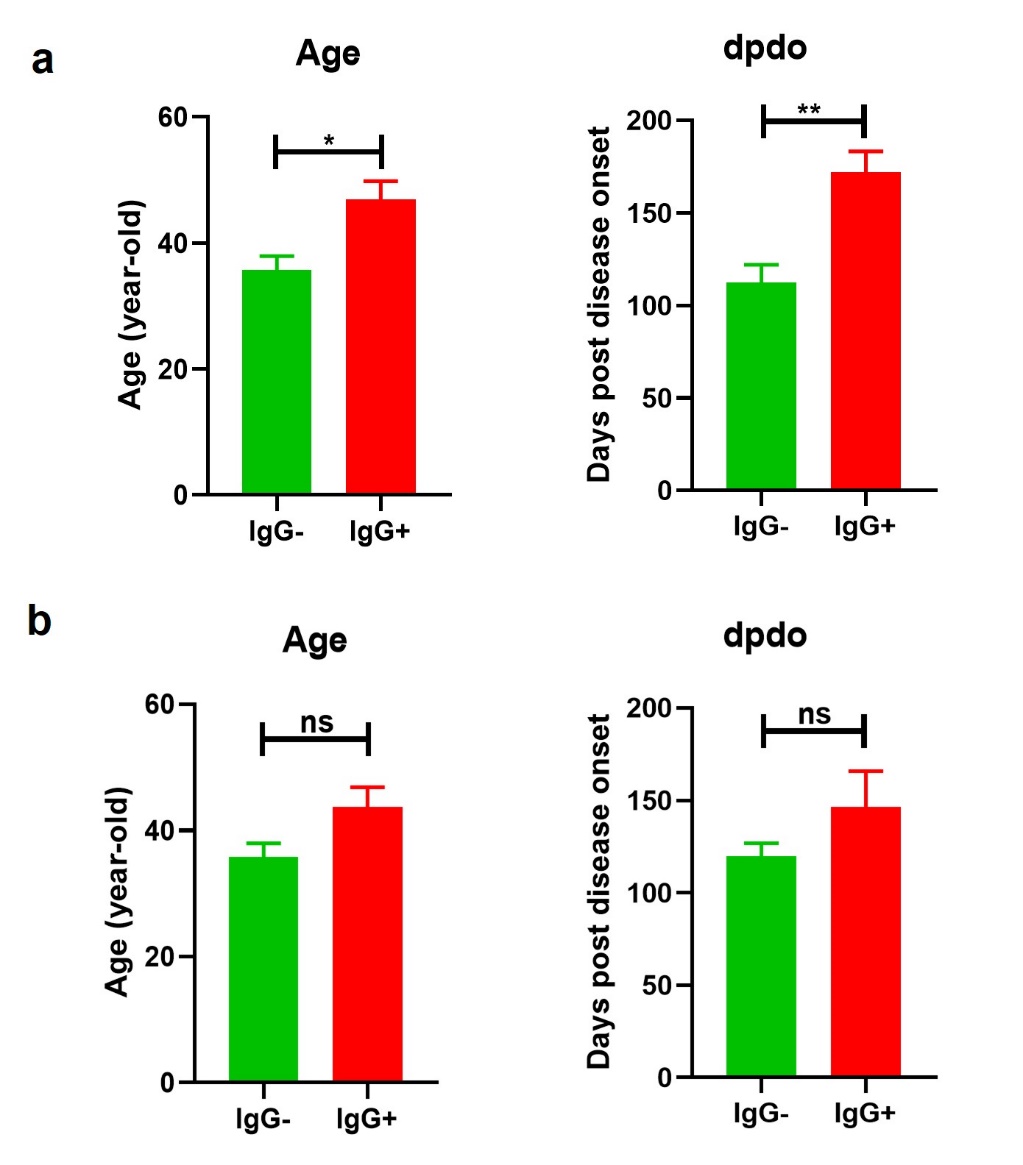

**Figure S6. Comparison of age and time following disease onset between IgG-seronegative and IgG-seropositive COVID-19 convalescent individuals.** (a) The age and days post disease onset (dpdo) are compared between IgG-seronegative (IgG-, n=8) and all IgG-seropositive (IgG+, n=23) CIs. (b) The age and days post disease onset (dpdo) are compared between IgG-seronegative (IgG-, n=8) and 7 selected IgG-seropositive (IgG+, n=7) CIs. Statistically significant differences are indicated by asterisks (* < 0.05, **< 0.01, Non-parametric Mann-Whitney test).
